# Attention and reinforcement learning in Parkinson’s disease

**DOI:** 10.1101/2020.09.12.294702

**Authors:** Brónagh McCoy, Rebecca P. Lawson, Jan Theeuwes

**Affiliations:** Department of Psychology, University of Cambridge, UK; Department of Experimental and Applied Psychology, Vrije Universiteit Amsterdam, The Netherlands; Institute for Brain and Behavior Amsterdam (iBBA), The Netherlands

## Abstract

Dopamine is known to be involved in several important cognitive processes, most notably in learning from rewards and in the ability to attend to task-relevant aspects of the environment. Both of these features of dopaminergic signalling have been studied separately in research involving Parkinson’s disease (PD) patients, who exhibit diminished levels of dopamine. Here, we tie together some of the commonalities in the effects of dopamine on these aspects of cognition by having PD patients (ON and OFF dopaminergic medication) and healthy controls (HCs) perform two tasks that probe these processes. Within-patient behavioural measures of distractibility, from an attentional capture task, and learning performance, from a probabilistic classification reinforcement learning task, were included in one model to assess the role of distractibility during learning. Dopamine medication state and distractibility level were found to have an interactive effect on learning performance; less distractibility in PD ON was associated with higher accuracy during learning, and this was altered in PD OFF. Functional magnetic resonance imaging (fMRI) data acquired during the learning task furthermore allowed us to assess multivariate patterns of positive and negative outcomes in fronto-striatal and visual brain regions involved in both learning processes and the executive control of attention. Here, we demonstrate that while PD ON show a clearer distinction between outcomes than OFF in dorsolateral prefrontal cortex (DLPFC) and putamen, PD OFF show better distinction of activation patterns in visual regions that respond to the stimuli presented during the task. These results demonstrate that dopamine plays a key role in modulating the interaction between attention and learning at the level of both behaviour and activation patterns in the brain.

## INTRODUCTION

Dopaminergic effects on cognition in Parkinson’s disease (PD) have typically been investigated in two separate domains that probe different aspects of cognition: 1) changes in learning from feedback, e.g. probabilistic classification [1,2] or reversal learning tasks [3], and 2) changes in the ability to remain goal-focused and resist task-irrelevant information, e.g. attentional switching [4, 5, 6] or interference tasks [7]. These cognitive domains are known to depend on neural processing across several overlapping brain regions; most notably the dorsolateral prefrontal cortex (DLPFC) which is central to maintaining focused attention [6, 8, 9] and supports learning by holding associative relationships among events and recent reinforcements in working memory [10, 11, 12]. Here, we seek to develop a better understanding of the commonalities across these two domains in terms of dopaminergic processing by administering both *attentional* and *learning* tasks to healthy controls and PD patients (on and off dopaminergic medication).

Dopaminergic nuclei in the midbrain fire differentially depending on the positive or negative valence of an outcome, with firing magnitude representing the size of a *prediction error*, i.e. the difference between the expected value of a chosen stimulus and the outcome received for that action [13, 14]. These signals are transmitted to widespread brain regions via several dopaminergic pathways, including a nigrostriatal projection from the midbrain’s substantia nigra pars compacta (SNc) to the striatum, which is involved in reward processing [15], motivation [16], and movement [17,18]. The SNc also projects to several parts of the frontal cortex [19], particularly the DLPFC. DLPFC is strongly implicated in cognitive control, including the control of attention [20], inhibition of actions or thoughts [21,22], rule learning [5], and working memory processes [23].

The central role of DLPFC in cognitive control underlies our ability to stay focused and resist distracting stimuli or events (see 19 for a review). Dopaminergic modulation of the DLPFC is proposed to improve distractor resistance by stabilizing task-relevant, working-memory representations in the DLPFC and making them less vulnerable to new inputs [6, 24, 25]. DLPFC lesions in humans, as well as ablation of analogous PFC regions in rhesus monkeys, were found to be accompanied by reduced distractor resistance [8, 26]. Studies on visual selective attention also suggest that top-down modulation from the DLPFC plays a key role in sharpening feature- or object-based representations in visual regions [27,28]. For example, impaired dopamine-related distractor resistance has been associated with reduced connectivity between DLPFC and the visual regions important for encoding task-relevant stimuli [6]. DLPFC activation, as well as the connectivity between DLPFC and fusiform face area, was found to be perturbed after face compared to scene distracters [29]. Thus, the fronto-visual processing that is integral to maintaining visual attention, at the expense of distractibility, is therefore likely to depend on dopaminergic mechanisms.

Rewards are well-known to boost the representation of visual stimuli [30,31], although the unique influences of attention and reward and their interactions on visual processing have proven difficult to tease apart [32]. Since dopamine signalling underscores reinforcement learning, research has begun to investigate the influence of dopamine on visual processing. For example, dopamine decreases blood oxygen level dependent (BOLD) activity [33], and modulates the representation of rewarded features, in visual cortex [34]. Furthermore, univariate BOLD activity has been shown to decrease within the representation of a high compared to low reward-predicting cue in visual cortex [34]. This suggests that dopamine-driven prediction error signals may tag representations of highly rewarded features in visual cortex. It is therefore likely that, similar to the dopaminergic enhancement of visual attention, the representation of rewarded visual features is also enhanced by dopamine. This, however, has yet to be tested explicitly.

PD patients suffer from a depletion of dopaminergic neurons, leading not only to the motor deficits characteristic of the disease, but also to changes in cognitive functioning, such as learning from feedback [35, 36, 37]. Replacement dopamine medication in patients has proven beneficial for certain cognitive processes, such as attentional switching [4, 38], but dopamine-related impairments have been observed in learning contexts [35]. Much research suggests that these impairments are driven by reduced learning from negative outcomes when patients are on compared to off medication [1,35, 37, 39, 40], although other studies also report medication-related differences in learning from positive outcomes [41, 42, 43]. Overall, there is general consensus that increased dopamine levels by medication in PD leads to a positivity bias [37, 44, 45]. A recent study in PD patients showed that when off dopaminergic medication, patients showed greater distractibility than healthy controls [46]. This suggests that distractor-resistance is compromised by altered dopamine levels in PD. However, it is not clear how attention, or distractibility, impacts on learning from feedback and the extent to which these effects are dopamine dependent.

Here we consider how dopamine levels in PD: 1) influence the inhibition of task-irrelevant distractors by measuring the extent of distractor-resistance in a behavioural attentional capture task [47], 2) alter the role that these attentional effects may play during a reinforcement learning task (see also 37), and 3) affect neural representations of positive and negative outcomes during learning, by assessing BOLD activation patterns while participants performed a reinforcement learning task [2] in the MRI scanner. Our aim was to uncover commonalities in dopamine-related indices of distractibility and learning. We first measured levels of distractibility and assessed whether these play a dopamine-dependent role in reinforcement learning performance. We then used multivariate pattern analysis (MVPA) to address whether fronto-striatal and occipital regions, known to be involved in both learning and distractor inhibition, contain patterns that differentially code for positive and negative outcomes, and whether these patterns are altered by dopamine levels in PD.

## MATERIALS AND METHODS

### Participants

Participants in this study were recruited as part of a larger project on reinforcement learning in PD, with original results published elsewhere (see 37). 24 patients with PD were tested twice, once while ON their standard schedule of dopaminergic medication and once in a clinically-defined OFF state (>12 hours withdrawal). 24 age-matched healthy controls (HCs) were tested once. All patients were diagnosed by a neurologist as having idiopathic PD according to the UK Parkinson’s Disease Society Brain Bank criteria. This study was approved by the Medical Ethical Review committee (METc) of the VU Medical Center, Amsterdam. All participants provided written informed consent in accordance with the Declaration of Helsinki. See Supplementary Materials for information on participant exclusions.

### Tasks

Participants completed two experimental tasks: a behavioural visual attentional capture task (based on 47) outside the MRI scanner, and a probabilistic reinforcement learning task (based on [2]) while in the scanner.

#### Attentional capture task

The standard stimulus display contained several green circles and one green diamond, placed on an imaginary circle equidistant from central fixation (see Fig. 1a). The total number of stimuli (“set size”) was either five (small set size), seven (medium set size) or nine (large set size). In 50% of the trials, one of the circle stimuli was a red distractor. Each shape contained a white line element that could be oriented either vertically or horizontally with equal probability across trials and conditions. Participants were instructed to respond as fast as possible to the orientation of the line inside the diamond, by pressing the “z” key when the line was vertical and the “/” key when the line was horizontal. There were 120 trials per set size across five short experimental blocks. See Supplementary Materials for further task details.

**Fig. 1.**
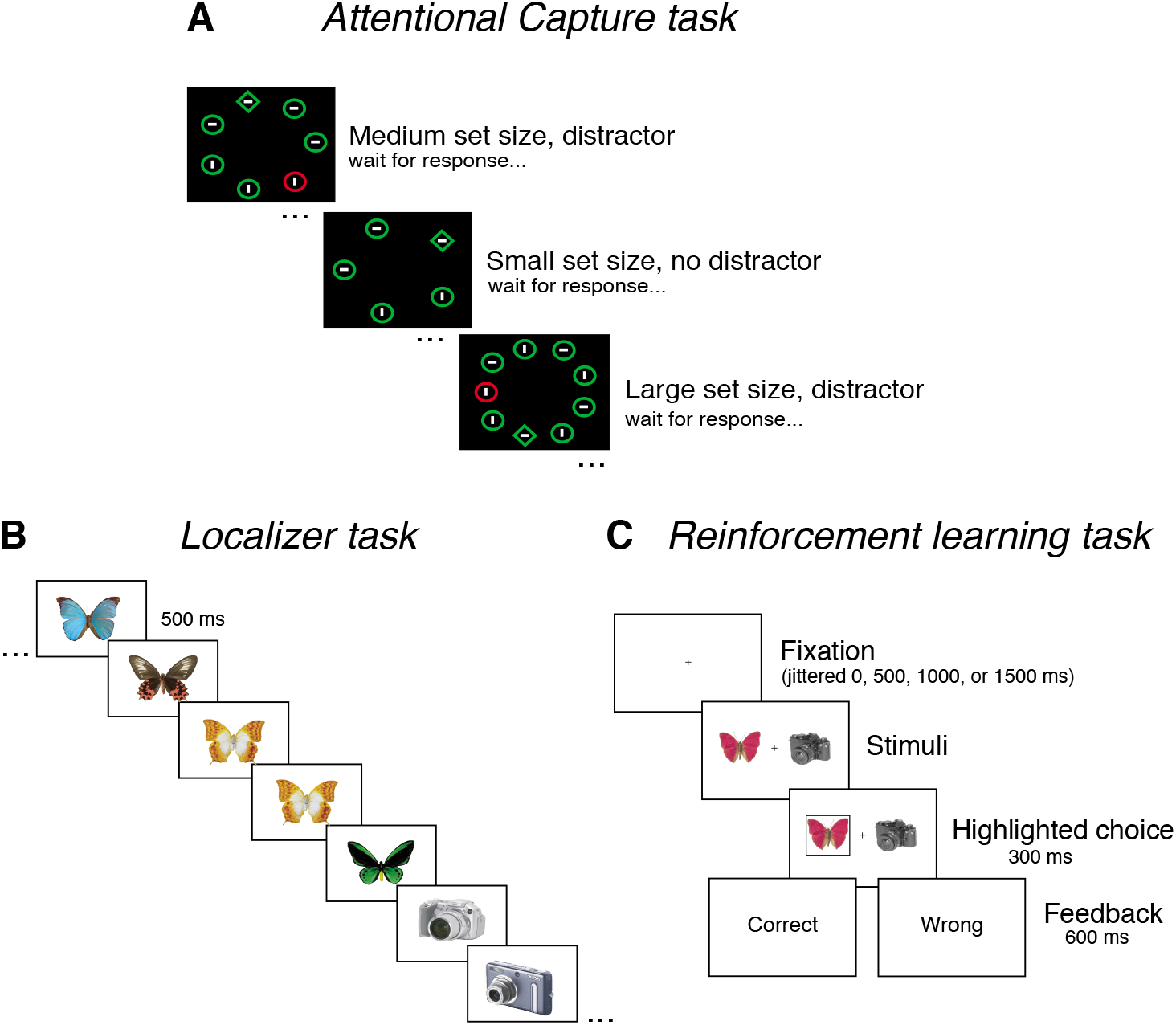
Experimental tasks. **(A)** A behavioural attentional capture task was performed outside the MRI scanner, to obtain a measure of distractibility, i.e. the amount of slowing down in no-distractor compared to distractor trials. **(B)** In the MRI scanner, participants first performed a localizer task, during which they pressed a button if two of exactly the same stimuli were presented sequentially. Stimuli consisted of many pictures from the same object categories as were used in the subsequent reinforcement learning task, as well as pink noise stimuli presented within the same central region as the object stimuli. A statistical contrast between object and noise trials provided voxels more involved in object-processing. Visual ROI masks were created on an individual subject level using the overlap of these voxels with the lateral occipital complex (LOC), and were labelled object-selective cortex (OSC). **(C)** Reinforcement learning task, also performed in MRI scanner (picture adapted from 37). Participants learned to choose the better option of each of three fixed stimulus pairings. Outcomes were probabilistic with a higher chance of receiving reward for one stimulus compared to the other; reward contingency was 80:20 for the easiest “AB” pair, 70:30 for the “CD” pair, 60:40 for the hardest “EF” pair.

#### Reinforcement learning task

In the learning phase, three different pairs of everyday object stimuli, e.g. shoes, balls, leaves (denoted as AB, CD and EF) were presented in random order. For each pair specific reward probabilities were associated with the individual stimuli, and participants had to learn to choose the best option of each pair based on the positive or negative feedback provided (see Fig. 1b). This contingency was 80:20 for the AB pair, indicating that the A stimulus would be rewarded 80% of the time it was chosen, whereas the B stimulus would be rewarded only 20% of the time. The reward probabilities were 70:30 and 60:40 for the CD and EF pairs, respectively. See Supplementary Materials for further details.

### Behavioural Analyses

#### Attentional capture task

Distractibility RTs (RT distractor absent - distractor present conditions) and absolute RTs for correct trials from the AC task were analyzed using repeated-measures ANOVA in JASP [48] and linear mixed-effects regressions with the *lme4* package in R [49, 50]. Repeated-measures ANOVAs were carried out separately per group comparison, either as within-subject comparisons for PD ON vs. OFF or between-group comparisons for PD ON/OFF vs. HC. See Supplemental Materials for further details and Supplementary Eq. 1 for the equation describing the linear mixed-effects regressions.

#### Reinforcement learning and distractibility

To quantify the distinct contribution of distractibility on learning, we carried out a mixed effects logistic regression on all trial-by-trial learning data. A similar analysis has previously been carried out on this data [37], however here we also include distractibility, along with stimulus pair (AB, CD, or EF), medication and disease status, to address our specific research questions (see Supplementary Eq. 2). The dependent variable encoded whether the better option of the stimulus pair was chosen on each trial. See Supplementary Materials for further details.

### fMRI Data Acquisition

Functional data for the reinforcement learning and localizer tasks were acquired using T2*-weighted echo-planar images with BOLD contrasts. See Supplementary Materials for full details.

### fMRI Localizer Analysis

Each fMRI scanning session began with a localizer run to extract a visual region of interest (ROI) for the MVPA analysis (see below). Stimuli in this run were objects of the same categories that were presented in the subsequent reinforcement learning task (six object categories per session), obtained from the same object database as the task stimuli [51]. The actual stimuli used in the learning task were not included. An object-selective cortex (OSC) ROI was taken as those voxels that responded more strongly to objects vs. noise, within anatomical masks of the superior and inferior lateral occipital cortex (LOC) (from the Harvard-Oxford Cortical Structural Atlas of the FSL package). See Supplementary Materials for full details.

### fMRI Single-trial Analysis

Single-trial whole-brain GLM analyses were performed on each participant’s reinforcement learning fMRI data to obtain trial-by-trial brain activation patterns. The single-trial regressors were locked to the onset of positive/negative feedback presented at the end of each trial. The analyses were carried out using Nipype’s FSL interface [52]. Full details are provided elsewhere (see Supplementary Materials in 37). One notable difference between the analysis described here and that described in [37] is that data were unsmoothed for the current MVPA procedure. Unsmoothed data were used because MVPA analyses require as fine-grained information as possible to more accurately classify brain representations of interest [53]. These GLM analyses resulted in one contrast of parameter estimates (COPE) for each trial of every participant and scanning session.

### Regions of Interest (ROIs)

In addition to the OSC ROI described above (created using functional localiser data), we created anatomical masks for four other relevant frontal and striatal ROIs using the Harvard-Oxford Cortical Structural Atlas in FSL; DLPFC (middle frontal gyrus), caudate nucleus, putamen and nucleus accumbens.

### fMRI Multivariate Pattern Analysis

A supervised machine learning technique using MVPA was carried out with the Python-based *scikit-learn* package [54]. Classification of events was done on a per run basis, i.e. across each learning run separately, and classification results were averaged across runs per participant. The MVPA analysis allowed us to identify patterns in the brain that can distinguish between positive and negative feedback. To obtain group results per ROI, classification accuracies and number of samples per participant were entered into a univariate normal-binomial model with a variational Bayes implementation using the MICP package in the TAPAS Matlab toolkit [55, 56]. See Supplementary Materials for full details.

## RESULTS

### Attentional capture: distractibility by task-irrelevant stimuli

We first analysed RTs from the AC task, for correct trials only (see *Methods*). Absolute RTs per set size, group and distractor condition are provided in Table 1. Statistical comparisons were made on a derivative of these quantities; distractibility RT, which was calculated as the within-participant difference between the distractor and no distractor conditions (within each of the ON and OFF medication sessions in PD patients). A higher value means that a participant took longer to respond to the target when a distractor was present (see Fig. 2a).

**Table 1:** RTs in the attentional capture task. Absolute RTs are shown for correct trials only. Mean and standard deviation (std) are provided across stimulus set size, group, and distractor conditions.

| *Set size / Group* | **HC** | **PD ON** | **PD OFF** |
| --- | --- | --- | --- |
| **Small** |  |  |  |
| Distractor absent (mean ± std) | 1005.5 ± 179.7 ms | 1150.0 ± 254.8 ms | 1166.4 ± 294.8 ms |
| Distractor present (mean ± std) | 1011.6 ± 165.7 ms | 1160.4 ± 278.3 ms | 1192.2 ± 288.9 ms |
| **Medium** |  |  |  |
| Distractor absent (mean ± std) | 1033.9 ± 159.0 ms | 1182.8 ± 272.7 ms | 1186.8 ± 299.5 ms |
| Distractor present (mean ± std) | 1070.3 ± 185.1 ms | 1215.4 ± 272.8 ms | 1238.0 ± 289.8 ms |
| **Large** |  |  |  |
| Distractor absent (mean ± std) | 1064.4 ± 176.7 ms | 1236.5 ± 282.6 ms | 1241.8 ± 317.8 ms |
| Distractor present (mean ± std) | 1102.2 ± 185.8 ms | 1244.0 ± 263.8 ms | 1308.8 ± 318.6 ms |

**Fig. 2.**
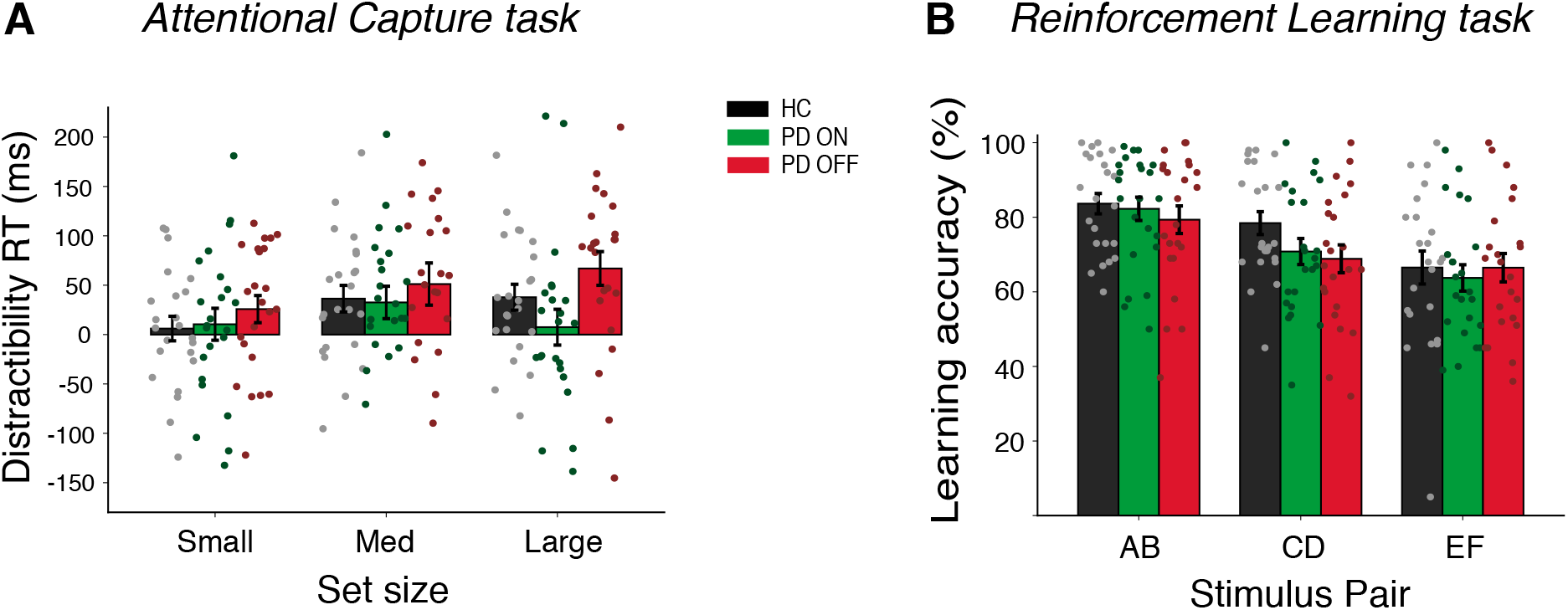
Behavioural results. **A)** *Attentional capture*: RT as a measure of attentional capture (“resistance” to distraction), per HC, PD ON and PD OFF group. RT distractibility is the amount of slowing caused when the distractor is present. **B)** *Reinforcement learning*: learning accuracy, i.e. the percentage of trials for each stimulus pair in which participants chose the ‘better’ option of the pair, per group. This part of the figure is adapted from 37.

We carried out separate repeated-measures ANOVAs on these distractibility RTs, per group comparison, e.g. PD OFF vs. ON, PD OFF vs. HC, and PD ON vs. HC. For the within-subject PD OFF vs. ON comparison, medication status, set size, and their interaction were included as independent variables. We found a main effect of medication on distractibility (F(1,22)=8.53, p=.008). This suggests that PD OFF medication were more distractible than PD ON. There was no evidence for a main effect of set size, nor a set size * medication interaction effect (both p>.1). For the PD OFF/ON vs. HC comparisons, there was tentative evidence for a main effect of set size for PD OFF vs. HC only (F(2,88) = 2.96, *p*=.057). There was no effect of group and no set size * group interaction for either comparison (all p>.09).

Next, a linear mixed-effects regression analysis was carried out on correct trial-by-trial log-transformed RTs (to account for positive skewing of the raw RT distributions) across all participants, to address both fixed experimental effects and random within-subject effects within one model (see Eq.1). We found main effects of distractor (β = .028, SE = .009, t = 3.26, *p* = 0.001) and set size (β = .004, SE = .005, t = 6.86, *p* <.001). There was a group effect of disease, i.e., HC vs. PD OFF (β = −.012, SE = .006, t = −2.21, *p*=0.032). Finally, there was also a significant interaction between medication status and distractor condition (β = −.022, SE = .009, t = −2.48, *p* = 0.013). This indicates that PD ON had less AC than OFF when distracting stimuli were present. We found no other significant effects.

### Role of distractibility in reinforcement learning

Next we characterised how distractibility affects reinforcement learning performance. We ran a linear mixed-effects regression on trial-by-trial data from the reinforcement learning task, with accuracy in choosing the better option of each pair as the dependent variable, and mean distractibility across all set sizes per participant included as an independent variable. See *Methods* and Supplementary Materials for a full description of the model, which includes disease and medication status as binary covariates (see Eq. 2 and 37). Overall accuracies in choosing the best option per stimulus pair and group can be seen in Fig. 2b. As detailed elsewhere (see 37), we found a main effect of stimulus pair (β = 0.217, SE = 0.078, z = 2.78, Pr(>|z|) = 0.005), and medication status (β = 0.489, SE =0.113, z = 4.31, Pr(>|z|)<.001), and both a stimulus pair * medication interaction (β =0.605, SE = 0.120, z = 5.06 Pr(>|z|)<001), and stimulus pair * disease interaction (β =.577, SE =.140, z=4.13, Pr(>|z|)<.001). Importantly for the current study, we found an interaction effect of medication status and mean distractibility RT on choice accuracy during learning (β = −0.606, SE = 0.187, z=-3.24, Pr(>|z|) = 0.001). The negative β-estimate here indicates that PD ON patients who showed less distractibility during the AC task were more accurate during the reinforcement learning task. Furthermore, results showed a 3way interaction effect of medication status, mean distractibility RT and stimulus pair (β = −0.700, SE = 0.190, z=-3.68, Pr(>|z|) < .001), and a 3-way interaction effect of disease, mean distractibility RT and stimulus pair (β = −0.547, stderr=0.206, z=-2.66, Pr(>|z|) = 0.008). To probe these 3-way interactions, we carried out separate mixed-effects regression models per group to assess whether the distractibility RT*stimulus pair interaction was present for all groups (Fig. 3). We found a significant distractibility RT*stimulus pair interaction for the HC group (β = −0.366, SE = 0.162, z = −2.26, *p* = 0.024) and PD ON group (β = −0.50299, SE = 0.14161, z = −3.55, *p* > .001) only. Analysis of the PD OFF data did not show any evidence for this interaction (*p*>.1). This indicates that the lesser the degree of distractibility in HC and PD ON participants, the better they performed at the easiest relative to most difficult stimulus pair choice during learning. Lower dopamine levels in PD OFF appear to alter this relationship.

**Fig. 3.**
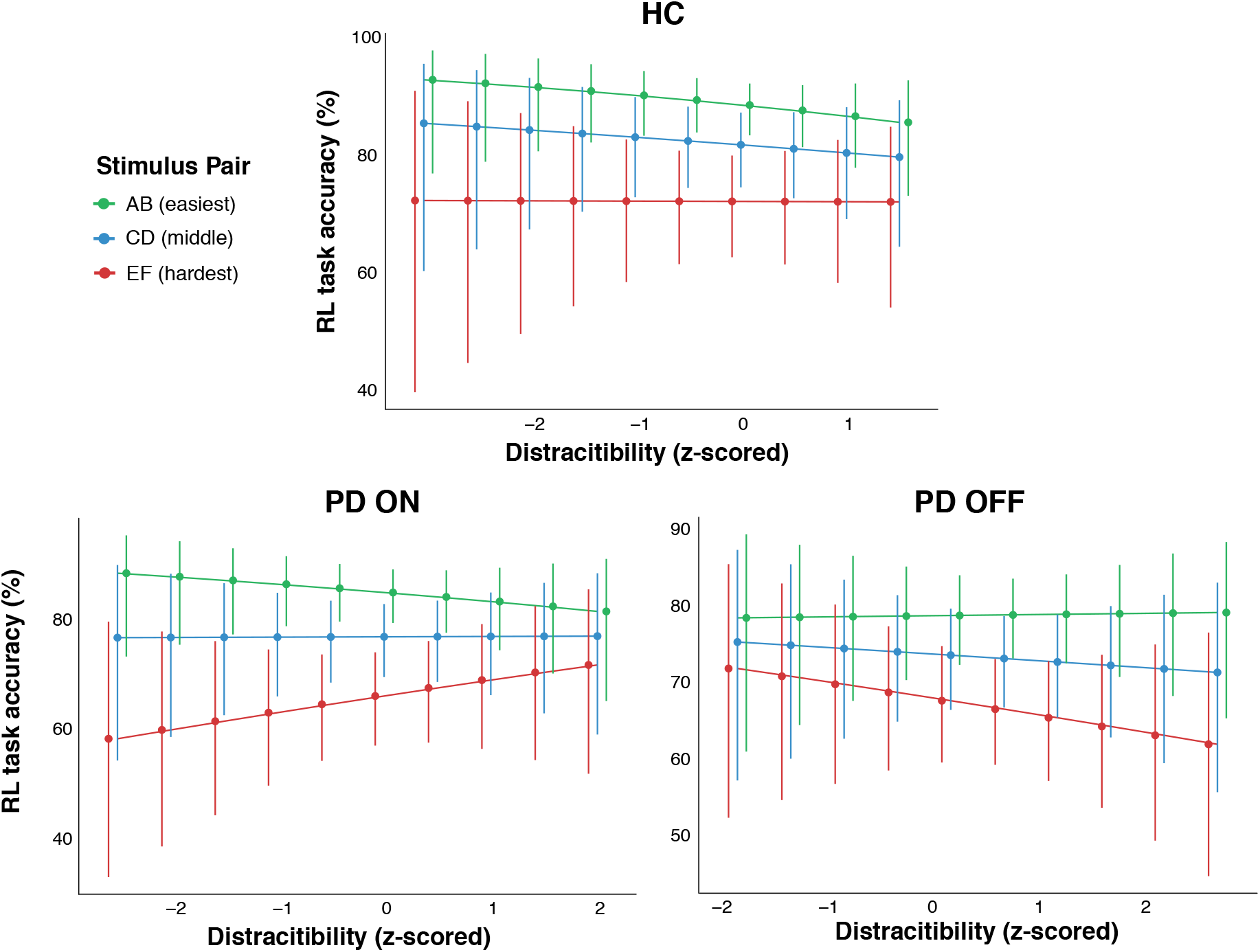
Effects of distractibility on learning. Interaction between level of distractibility (z-scored) from the AC task and stimulus pair (easy to difficult) on choice accuracy during the RL task, for HC, PD ON and PD OFF groups. Different patterns between PD OFF and each of the HC and PD ON groups represent the significant interaction effects of these variables. Errorbars represent 95% confidence intervals.

### fMRI multivariate pattern analysis of positive vs. negative outcomes during learning

MVPA of each participant’s trial-by-trial BOLD data was carried out in *a priori* ROIs (see Methods and Fig. 4). These ROIs were: visual OSC (defined using functional localizer data), and the fronto-striatal regions DLPFC, caudate nucleus, putamen and nucleus accumbens (using anatomically-defined masks). In all ROIs and groups, classification accuracies were significant for positive vs. negative outcomes (all *p*<.001). See **Fig. 4** for classification accuracies per group and ROI.

**Fig. 4.**
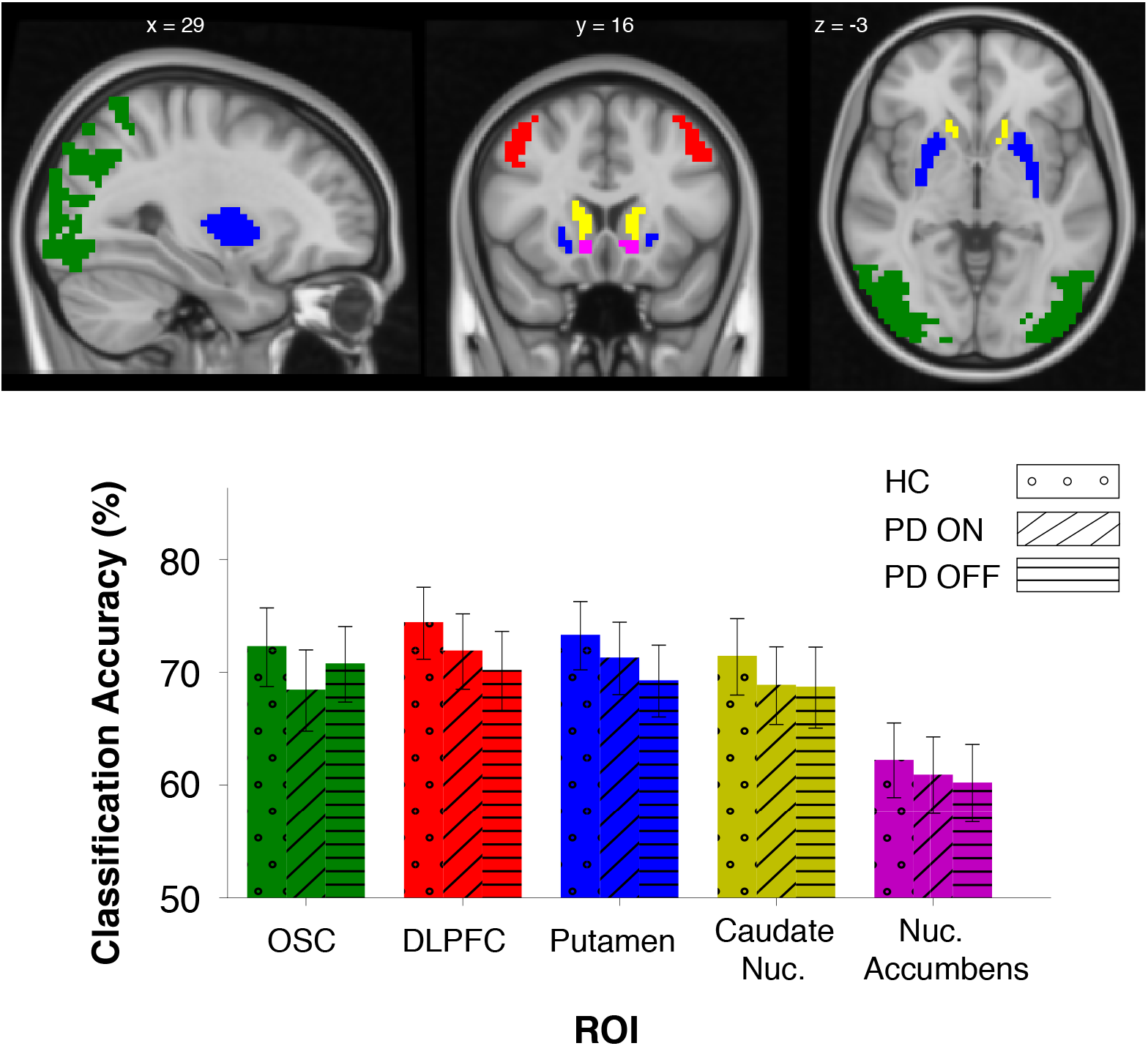
Classification accuracy results per group and ROI. Functional and anatomical masks are displayed as follows: OSC (green), DLPFC (red), putamen (blue), caudate nucleus (yellow), nucleus accumbens (magenta). The OSC was based on a functional localizer (see Methods) and was different per participant; mask shown here is from one sample participant. All other masks were anatomically-defined and were applied to all participants. Classification accuracy results presented are as follows; ***OSC***: HC (72.31% [68.73, 75.70]), PD ON (68.46% [64.78, 71.98]), PD OFF (70.78% [67.35, 74.05]); ***DLPFC***: HC (74.44% [71.15, 77.53]), PD ON (71.92% [68.49, 75.17]), PD OFF ( 70.2% [66.61, 73.62]); ***Putamen:*** HC (73.32% [70.21, 76.26]), PD ON (71.3% [68.01, 74.44]), PD OFF (69.29% [66.04, 72.40]); ***Caudate Nucleus***: HC (71.45% [67.98, 74.75]), PD ON (68.89% [65.37, 72.25]), PD OFF (68.72% [65.05, 72.23]); ***Nucleus Accumbens:*** HC (62.23% [58.88, 65.50]), PD ON (60.93% [57.51, 64.27]), PD OFF (60.22% [56.77, 63.60]).

In OSC, classification accuracy in HC was significantly higher than in PD ON (t(40) = 2.14, *p* = .038). The within-PD subject comparison showed greater accuracies in PD OFF than ON, which was not statistically significant (t(20) = 1.93, *p*=.068). There were no significant differences in HC vs. PD OFF (*p*>.1).

In DLPFC, we found significantly higher decoding accuracy in HC compared to PD OFF (t(42)=2.50, p=.017). There were no significant differences between HC and PD ON or in PD patients ON vs. OFF medication (both *p*>.1).

In the putamen, we also found significantly higher classification accuracies in HC compared to PD OFF (t(43)=2.94, *p*=.005). There were no differences between HC and PD ON nor between PD ON and OFF (both *p*>.1). Statistical comparisons in the caudate nucleus and nucleus accumbens showed no pair-wise significant differences between any of the HC, PD ON, and PD OFF groups.

We next calculated within-PD patient differences in classification accuracy in each of these ROIs and carried out paired t-tests between fronto-striatal ROIs and the visual OSC ROI. Previous research has indicated a dopaminergic role in top-down control of visual regions [28]. Here, we sought to explore how relative information, i.e. patterns of activation, in these regions may differ according to dopamine levels (see Fig. 5). We found a significantly higher PD ON-OFF difference in classification accuracy both in DLPFC compared to OSC (t(20) = 2.99, *p* = .007; DLPFC: 1.72%± 1.77% SEM; OSC: −2.31%±1.71% SEM) and in the putamen compared to OSC (t(20) = 3.01, *p* = .006; Putamen: 1.76%±1.67 SEM). There was no significant dopamine-related difference between the caudate nucleus and OSC, nor between the nucleus accumbens and OSC.

**Fig. 5.**
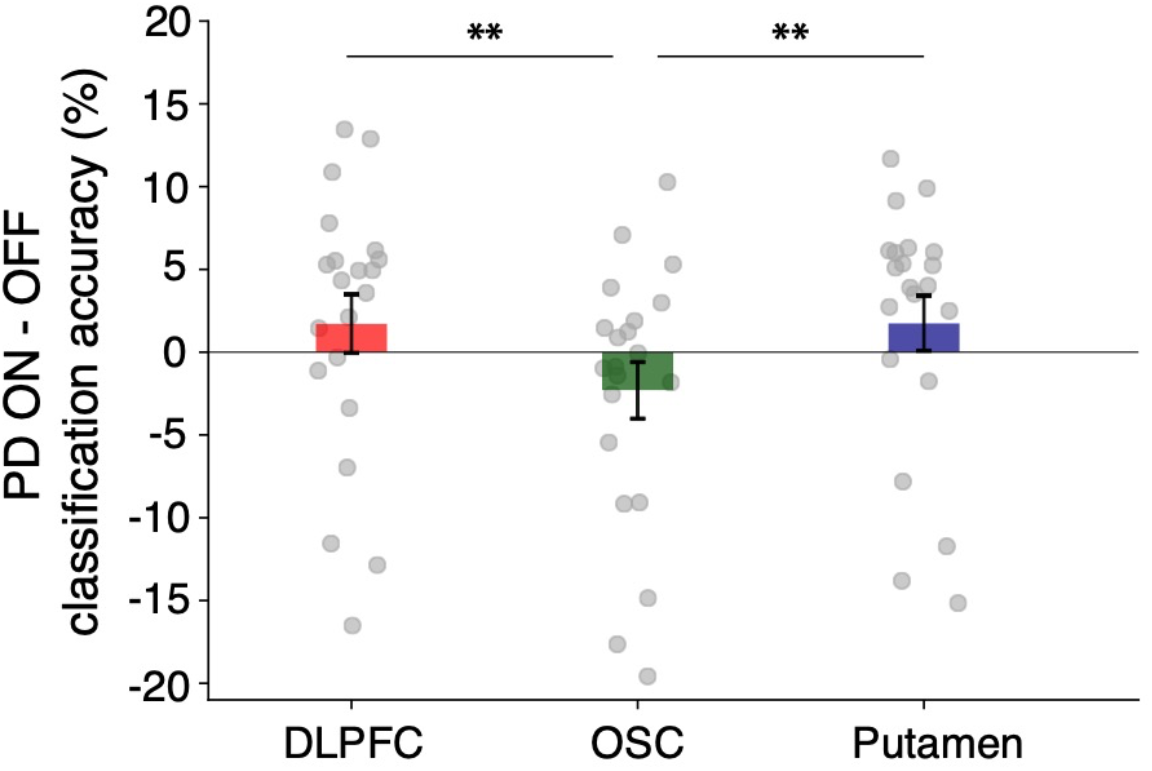
Within-PD patient ON vs. OFF classification accuracy differences in the DLPFC, OSC, and putamen. Error bars are ± 1 SEM. ** represents significant differences between regions at *p*<.01.

## DISCUSSION

In this study we report on several behavioural and brain mechanisms claimed to be altered in PD, by means of the disease itself or by dopaminergic medication used to treat associated motor symptoms. PD patients and HCs performed an attentional capture task, to assess levels of distractibility by task-irrelevant items in the environment, and a reinforcement learning task, to describe how these groups learn from the outcomes of actions.

Our behavioural results show that PD ON demonstrate a significantly greater ability to resist distracting stimuli than PD OFF, as indicated by reduced RT interference in the distractor vs. no distractor condition in the AC task. We furthermore found that medication status and mean distractibility RT interact to affect choice accuracy during learning, specifically; PD ON patients who showed less distractibility in the AC task performed better in the reinforcement learning task.

MVPA of functional neuroimaging data from the reinforcement learning task shows a medication-related effect in representations that distinguish positive from negative feedback across fronto-striatal and visual brain regions. Specifically, we found a medication-related interaction between DLPFC and OSC, with greater classification accuracy in PD ON (vs. OFF) in DLPFC, but greater accuracy in PD OFF (vs. ON) in OSC (see Fig. 4). A similar interaction was found between the putamen and OSC. This suggests that brain activations in these fronto-striatal regions are more separable for positive compared to negative outcomes when PD patients are ON medication, whereas visual brain regions show more separable outcome responses when patients are OFF medication. Higher classification accuracies were furthermore found in HC participants compared to PD OFF in DLPFC and the putamen, and compared to PD ON in OSC. These results compliment the within-patient findings shown in Fig. 4.

Our finding that dopaminergic medication improves the ability to resist task-irrelevant (salient) stimuli in the environment dovetails with theory and results from previous research describing the role of dopamine in distractor inhibition [6, 24, 25, 46, 57, 58]. These studies suggest that dopaminergic effects on working memory, via the excitatory D1 and inhibitory D2 pathways in the basal ganglia and PFC, relate to improvements and impairments in the ability to resist distracting input, respectively [6]. Prior research of ours shows that reward magnitude plays a role in distractor inhibition, with an increase in erroneous saccades to distractors that signal increasing levels of reward in the environment [59]. This highlights the interplay between (irrelevant) reward signals and the ability to remain goal-directed. Dopamine in fronto-striatal regions is strongly implicated in rewarding and motivational responses [60]. Although the distractor stimulus in the current AC task was not associated with reward, the dopamine-related improvements in PD patients show a shift in this balance of AC towards task-irrelevant stimuli vs. task-relevant goals, that is likely mediated by these fronto-striatal regions.

The relationship between AC and learning from rewards or reinforcements is further suggested by our findings regarding the combined influence of distractibility (as indexed by overall RT differences between distractor conditions) and dopaminergic medication on performance accuracy during learning. Investigating how the balance between distractibility and goaldirectness in PD patients ON vs. OFF medication may shift in accordance with varying distractor reward magnitude in the AC task is an interesting avenue for future research.

Our MVPA findings align with the suggestion of medication-related changes in the balance between distractibility and goal-directedness. We show that positive and negative outcomes can be more easily distinguished in PD ON than OFF in two fronto-striatal regions, the DLPFC and putamen, whereas these events are more easily decoded in PD OFF than ON in the visual OSC. This provides some evidence for PD ON relying more on information from higher-level control- and reward-related regions during learning, with PD OFF being more visually guided by lower-level, salience-related regions. This notion of competition between top-down vs. bottom-up processing has been long discussed in the literature [61, 62, 63, 64, 65]. OSC functional masks in this study were created on an individual basis, using a functional localizer to create an objects - noise contrast. Although this focuses on visual object-related regions, rather than feedback-related regions, positive and negative outcomes can still be easily classified. Several previous studies have shown reward-related signals in visual cortex [30, 31]. According to reinforcement learning theory, at the time of the outcome, positive or negative feedback is used to update the value of the chosen object [66, 67]. This presumably requires the integration of the object representation itself with the outcome and/or new object value. On the basis of our results we speculate that reward binding to objects may occur more at a stimulus-driven, visual level in PD OFF but more at a conceptual, reward-driven, and working-memory level in fronto-striatal regions in PD ON. It has been shown that disrupted dopamine-related distractor resistance is associated with reduced connectivity between DLPFC and the visual regions recruited for encoding task-relevant stimuli [6]. Interestingly, dopamine has previously been associated with a reduction of univariate activation in visual regions [33]. Similarly, reward delivery without visual stimulation leads to decreased univariate activity within reward-associated cue representations in visual cortex [34]. Although it is not possible to make direct comparisons between these studies and our MVPA results, the current findings extend the notion that dopamine and reward lead to differential activations in visual regions. It is possible that higher dopamine levels in PD ON reduce univariate activity within the representation of positive outcomes compared to PD OFF, thereby interfering with multivariate patterns that may more easily distinguish between positive and negative outcomes.

A limitation of the current study relates to the flexibility of MVPA. Due to the simultaneous presentation of two stimuli on each trial, as well as extremely large differences in choosing the better vs. worse option of pairs, e.g. some participants may only have chosen the worst “B” option only several times across the whole experiment, it was not possible to appropriately tease out individual stimulus representations for pattern classification. An examination of dopamine-related differences in DLPFC representations of individual objects and how these might relate to distractibility, e.g. by presenting the objects as distractor stimuli in a separate task, is a compelling avenue for future research.

In summary, the current study examines the role of dopamine in the interplay between attention and learning. We provide evidence for dopamine-related effects of attentional distractibility on reinforcement learning, as well as a dopamine-related dissociation of multivariate representations in executive control, reward-associated and stimulus-driven brain regions. This application of MVPA is rare in the field of PD research and, when used in conjunction with other modern fMRI pre-processing methods, highlights its future potential in distinguishing event- or stimulus-induced brain representations in PD.

## AUTHOR CONTRIBUTIONS

BM and JT conceived of the experiments. BM carried out data collection and data analysis. BM wrote the first draft of the manuscript. BM, RPL and JT contributed to the interim and final versions of the manuscript.

