## Supplementary Information for "Attention and reinforcement learning in Parkinson’s disease"

### Exclusions

Data from one PD patient and one HC were excluded from analyses, due to falling asleep in the scanner and insufficient learning of the reinforcement learning task (<60% accuracy across all stimulus pairs), respectively. The anatomical scan of one HC was furthermore not collected due to requested early termination of scanning. Full participant details, inclusion and exclusion criteria, and medication information is included elsewhere (see 1).

### Tasks

*Attentional capture task*

Participants performed the task on a laptop outside the scanning room. A warning beep sounded to inform participants when they made an error. The session consisted of one practice block of 72 trials and 5 experimental blocks of 72 trials. Each set size contained 24 trials per block, totalling 120 trials per set size across all experimental blocks. Participants pressed the space bar when they were ready to begin a new block. A trial started with the presentation of a central fixation cross for 1000ms, after which the stimulus display was shown. There was no time limit placed on responding to ensure that slowed motor-related responses of PD patients did not result in lost trials. Data were later analyzed with a threshold of 3000ms to ensure that responses due to inattention to the task were not included. This is a liberal threshold compared to other studies using this task, e.g. [2,3], which used a 1600ms threshold. However, those studies were in young, healthy participants. We therefore relaxed this threshold to allow for age- and disease-related slowing.

*Reinforcement learning task*

Participants were instructed to try to find the better option of a pair in order to maximize reward. Outcomes were either "Goed" or "Fout" text (meaning "correct" or "wrong" in Dutch), leading to a pay-out of 10 cents for correct trials and nothing for incorrect trials. Participants completed two learning runs, with 50 trials per stimulus pair per run. Object stimuli were obtained from an existing stimulus database [4].

**Behavioural Analyses**

*Attentional capture task*

Error rates were extremely low as participants could take as long as required to choose the correct response; these were therefore not analysed. The linear mixed-effects model incorporated both fixed and random trial-by-trial effects. RTs were log transformed to overcome positive skewing of the raw distribution and included as the dependent variable of the model. The fixed effects variables were distractor (absent/present), set size (small, medium, large) and their interaction. Two binary covariates were also included (as in 1 and 5); the between-subject effect of disease (*Dis*, where PD = 0, HC = 1) and the within-subject effect of dopaminergic medication state (*Med*, where OFF = 0, ON = 1), along with their interactions with the fixed effects variables. Participant was entered as a random effect [6]. Random slopes were included for distractor and set size to capture additional variability at the subject level [7]. The full model was therefore set up according to Eq. 1.

***(Eq.1)***

*log(RT) = Distractor + Setsize + Dis + Med + Distractor*Setsize + Distractor*Med + Distractor*Dis + Setsize*Med + Setsize*Dis + (1 | Participant) + (Distractor | Participant) + (Setsize | Participant)*

*Reinforcement learning and distractibility*

The mixed effects logistic regression analysis carried out on learning and distractibility data is robust for assessing within- and between-subject individual differences and has been previously been used to examine dopaminergic effects on learning [5, 8]. The dependent variable encoded whether the better option of the stimulus pair was chosen on each trial (correct = 1, incorrect = 0). The within-subject (random-effect) explanatory variable was stimulus pair (AB = 1, CD = 0, EF = -1). We also included two binary covariates, each interacting with stimulus pair: the between-subject effect of disease (*Dis*: Parkinson's disease (PD) = 0, control = 1), and the within-subject effect of dopaminergic medication (*Med*: OFF = 0, ON =1, with HCs considered as being in an OFF state). Distractibility RTs from the AC task, i.e., the mean difference between distractor present vs. distractor absent trials, were mean-centered across all participants and included in the model as a fully interacting covariate. PD OFF therefore acted as the baseline group (*Med*=0, *Dis*=0), which allowed comparison to PD ON (where the Distractibility_RT**Med* interaction reflects the interacting effect of medication and distractibility on learning) and to HC (where the Distractibility_RT **Dis* term reflects the interacting effect of disease (HC or PD OFF) and distractibility on learning). In addition to this model (see Eq. 2), similar separate models were carried out per HC, PD ON, and PD OFF groups, without the *Med* and *Dis* terms. Statistical analyses were performed in R [9] using the lme4 package [10].

***(Eq.2)***

*Correct = Stim_pair + Distractibility_RT + Med + Dis + Stim_pair*Distractibility_RT + Stim_pair*Med + Stim_pair*Dis + Distractibility_RT*Med + Distractibility_RT*Dis + Distractibility_RT*Med*Stim_pair + (1 | Participant)*

### fMRI Data Acquisition

BOLD fMRI data were acquired using a 3T GE Signa HDxT MRI scanner (General Electric, Milwaukee, WI, USA) with 8-channel head coil at the VU University Medical Center (Amsterdam, The Netherlands). Each brain volume contained 42 axial slices, with 3.3 mm in-plane resolution, TR = 2,150 ms, TE = 35 ms, FA = 80 degrees, FOV = 240 mm, 64 x 64 matrix. The first two TR volumes were removed from each run to allow for T1 equilibration. Structural images were acquired with a 3D T1-weighted magnetization prepared rapid gradient echo (MPRAGE) sequence with the following acquisition parameters: 1 mm isotropic resolution, 176 slices, repetition time (TR) = 8.2 ms, echo time (TE) = 3.2 ms, flip angle (FA) = 12 degrees, inversion time (TI) = 450 lms, 256 x 256 matrix. The subject's head was stabilized using foam pads to reduce motion artifacts. Preprocessing of fMRI data was carried out using FMRIPREP version 1.0.0-rc2 [11, 12], a Nipype-based tool [13]. Full details of the scanning protocol and preprocessing are included elsewhere (see 1). Below, we include details relevant to the specific analyses that targeted the current research questions.

### fMRI Localizer Analysis

Stimuli were centrally-presented within a black square frame kept constant in size across the run. Pink noise within these square patches were also included as stimuli. The localizer run consisted of four similarly-structured blocks. Each block was made up of eight mini-blocks, containing either stimuli from each of the six object categories that were used in the subsequent reinforcement learning task, pink noise, or a null mini-block during which only the central fixation cross was presented. In all but the null mini-block, stimuli were flashed briefly for 500 ms, with 300 ms fixation-cross only intervals. There were ten distinct object stimuli per mini-block, presented twice, making up 20 stimulus presentations per mini-block. Each mini-block lasted 16 seconds. This localizer run lasted ~ 9 min in total. Participants had to push a button when two consecutive object images were identical, to ensure they were paying attention to the stimuli. Data from the localizer run were corrupted in three participants (2 PD, 1 HC) and were excluded when creating functional OSC masks. Preprocessing was carried out as described in [1].

**fMRI Multivariate Pattern Analysis**

The unsmoothed fMRI data from the single-trial analysis was used as input and each trial was labelled as either 'good' or 'bad' feedback. Activity patterns ("features") were standardized per run by removing the mean across that run and scaling to unit variance. Since positive and negative feedback was provided probabilistically, depending on the presented stimulus pair and the stimulus chosen, it was not possible to balance the number of positive and negative feedback events, neither across run nor across participant. Overall, participants received more positive than negative feedback since their task was to learn which was the better stimulus of each pair. Such an imbalance in the number of positive vs. negative samples provided to the classifier can lead to a bias in the classification procedure and should be addressed. To account for this, we used a combination of oversampling and undersampling with the *SMOTETomek* function provided in the *imbalanced-learn* Python package [14]. Classification accuracy score was estimated per learning run and overall accuracy was averaged across runs.

This takes both fixed (within-subject) and random (between-subject) effects into account in one model. A hierarchical structure is employed by this model for mixed effects analysis, using a variational Bayes approach. Variational Bayes is more efficient (albeit, less exact) than more standard Markov chain Monte Carlo (MCMC) sampling procedures used for inference in Bayesian modelling. This model was set up separately per HC, PD ON and PD OFF group. Statistical comparisons between groups were performed using either paired-samples t-tests (PD ON vs. OFF) or independent-samples t-tests (HC vs. PD ON/OFF) on the original subject-level classification accuracies extracted using *scikit-learn*. To compare classification accuracies between fronto-striatal ROIs and the visual OSC ROI in PD patients ON and OFF medication, i.e. the relative fronto-striatal vs. visual involvement in classifying outcomes, we first found the within-ROI difference between ON/OFF sessions and compared these differences across ROIs. Due to exclusions in creating the OSC masks (see *Methods*), MVPA results from two participants were here dropped from the non-OSC ROIs to make the appropriate statistical comparisons across ROIs.
